# Ligand-Displaying *E. coli* Cells and Minicells for Programmable Delivery of Toxic Payloads via Type IV Secretion Systems

**DOI:** 10.1101/2023.08.11.553016

**Authors:** Yang Grace Li, Kouhei Kishida, Natsumi Ogawa-Kishida, Peter J. Christie

**Affiliations:** Department of Microbiology and Molecular Genetics, University of Texas Health Science Center, McGovern School of Medicine, Fannin St, Houston, Texas 77030; Graduate School of Life Sciences, Tohoku University, 2-1-1 Katahira, Aobaku, Sendai, 980-8577, Japan

**Author notes:** Correspondence: Peter J. Christie, Dept. Microbiology and Molecular Genetics, McGovern Medical School, 6431 Fannin St, Houston, TX 77030. **Author Contributions:** P.J.C. and Y.G.L. designed the research; Y.G.L, K.K., and N.O-K. performed the research; P.J.C, Y.G.L, K.K., and N.K. analyzed the data; P.J.C. and Y.G.L. wrote the paper with contributions from K.K. and N.K.

**Keywords:** conjugation, type IV secretion, CRISPR/Cas9, minicells, nanobodies, antimicrobials

## Abstract

Bacterial type IV secretion systems (T4SSs) are highly versatile macromolecular translocators and offer great potential for deployment as delivery systems for therapeutic intervention. One major T4SS subfamily, the conjugation machines, are well-adapted for delivery of DNA cargoes of interest to other bacteria or eukaryotic cells, but generally exhibit modest transfer frequencies and lack specificity for target cells. Here, we tested the efficacy of a surface-displayed nanobody/antigen (Nb/Ag) pairing system to enhance the conjugative transfer of IncN (pKM101), IncF (F/pOX38), or IncP (RP4) plasmids, or of mobilizable plasmids including those encoding CRISPR/Cas9 systems (pCrispr), to targeted recipient cells. *Escherichia coli* donors displaying Nb’s transferred plasmids to *E. coli* and *Pseudomonas aeruginosa* recipients displaying the cognate Ag’s at significantly higher frequencies than to recipients lacking Ag’s. Nb/Ag pairing functionally substituted for the surface adhesin activities of F-encoded TraN and pKM101-encoded Pep, although not conjugative pili or VirB5-like adhesins. Nb/Ag pairing further elevated the killing effects accompanying delivery of pCrispr plasmids to *E. coli* and *P. aeruginosa* transconjugants bearing CRISPR/Cas9 target sequences. Finally, we determined that anucleate *E. coli* minicells, which are clinically safer delivery vectors than intact cells, transferred self-transmissible and mobilizable plasmids to *E. coli* and *P. aeruginosa* cells. Minicell-mediated mobilization of pCrispr plasmids to *E. coli* recipients elicited significant killing of transconjugants, although Nb/Ag pairing did not enhance conjugation frequencies or killing. Together, our findings establish the potential for deployment of bacteria or minicells as Programmed Delivery Systems (PDSs) for suppression of targeted bacterial species in infection settings.

**IMPORTANCE:** The rapid emergence of drug-resistant bacteria and current low rate of antibiotic discovery emphasize an urgent need for alternative antibacterial strategies. We engineered *Escherichia coli* to conjugatively transfer plasmids to specific *E. coli* and *Pseudomonas aeruginosa* recipient cells through surface display of cognate nanobody/antigen (Nb/Ag) pairs. We further engineered mobilizable plasmids to carry CRISPR/Cas9 systems (pCrispr) for selective killing of recipient cells harboring CRISPR/Cas9 target sequences. In the assembled Programmed Delivery System (PDS), Nb-displaying *E. coli* donors with different conjugation systems and mobilizable pCrispr plasmids suppressed growth of Ag-displaying recipient cells to significantly greater extents than unpaired recipients. We also showed that anucleate minicells armed with conjugation machines and pCrispr plasmids were highly effective in killing of *E. coli* recipients. Together, our findings suggest that bacteria or minicells armed with PDSs may prove highly effective as an adjunct or alternative to antibiotics for antimicrobial intervention.

## INTRODUCTION

The programmable binding of bacteria or bacterial products such as minicells to other bacteria or eukaryotic cells has enormous potential for a range of basic (1), synthetic biology and biotechnological (2, 3), and cancer therapeutic applications (4–7). Such targeting systems rely on the surface display of a ligand on one bacterial cell that specifically binds a receptor displayed on a second cell. In early tests, Fos and Jun leucine zippers were surface-displayed on *Escherichia coli* through substitution of passenger domains of an IgA protease autotransporter. Upon mixing of strains displaying Fos or Jun, the leucine zippers dimerized and elicited strong intercellular aggregation (8). More recently, *E. coli* cells engineered to surface-display variable domains of camelid heavy-chain antibodies, termed nanobodies (Nb’s), were shown to selectively bind other bacterial or eukaryotic cell targets displaying cognate antigens (Ag’s) (6, 9).

In an era of increasing prevalence of antibiotic resistance, largely driven by the widespread use of antibiotics, there is an urgent need to develop alternative antimicrobial strategies. Ideally, such strategies fulfill two objectives. First, they target specific cell types, namely, pathogens as opposed to commensal species. Second, they effectively suppress growth or kill the targeted cells. Bacteriophage therapy is considered a promising alternative to antibiotics, as many phages selectively bind and infect only certain bacterial host species. Phage therapy, however, can be rendered ineffective by the rapid evolution of phage resistance or difficulties in identifying phages capable of infecting pathogens of interest (10, 11). Bacteria also have the capacity to target and kill other bacterial cells through the intercellular transfer of toxins via dedicated secretion systems (12). Recently, it was shown that the combined deployment of *Enterobacter cloacae* harboring one such killing machine designated as the type VI secretion system (T6SS), together with display of Nb’s on the cell surface, selectively killed *E. cloacae* or *E. coli* cells displaying cognate Ag’s (13). While an important advance, the applicability of T6SS-mediated killing is limited by the fact that many Gram-negative species that harbor T6SSs, including *E. cloacae,* are themselves opportunistic pathogens or emerging antibiotic resistance threats (14).

Here, we sought to deploy conjugation machines for the targeted delivery of plasmids encoding toxic elements. Conjugation machines, which comprise a large subfamily of the type IV secretion systems (T4SSs), mediate the widespread transmission of mobile genetic elements (MGEs) among most if not all species of bacteria (15, 16). We engineered *E. coli* donor cells with different model conjugation systems to produce Nb’s that bind cognate Ag’s displayed by recipient cells. The Nb-producing donors delivered conjugative or mobilizable plasmids to Ag-producing *E. coli* or *Pseudomonas aeruginosa* recipients at significantly higher levels than to unpaired recipients. Nb-producing donors also mobilized the transfer of plasmids encoding CRISPR/Cas9 (Clustered regularly interspaced short palindromic repeats/Cas9 nuclease) elements, resulting in killing of Ag-paired *E. coli* and *P. aeruginosa* recipients at significantly elevated frequencies relative to unpaired recipients. Remarkably, anucleate *E. coli* minicells also functioned as efficient donors of self-transmissible as well as toxic pCrispr plasmids to recipient cells, with the latter eliciting significant levels of killing. Our findings establish that Programmed Delivery Systems (PDSs), comprised of ligand-displaying bacterial cells or clinically-safer anucleate minicells with functional conjugation systems, may prove highly useful for targeted antimicrobial therapy.

## RESULTS

### Pairing of surface-displayed nanobodies and antigens promotes targeted pKM101 transfer

The IncN plasmid pKM101 is representative of a large number of conjugative plasmids that transfer at low or modest frequencies in conditions of dilute cell densities, e.g., liquid media, and at considerably higher frequencies under conditions of dense cellular growth, e.g., solid-surfaces or biofilms (17–19). To determine whether Nb/Ag pairing stimulates pKM101 transfer between *E. coli* cells, we tested four previously characterized Nb/Ag pairs, two identified in a high-throughput screen (Nb/Ag [X], Nb/Ag [Y]), one corresponding to an Nb that binds native intimin from enterohemorrhagic *E. coli* (EHEC) (Nb/Ag [Int]), and one that binds *E. coli* outer membrane protein BamA (Nb/Ag [BamA]) (9, 13). As null controls, donor or recipient cells produced the intimin β-barrel without an attached passenger domain (**Fig. 1A**).

**Fig 1.**
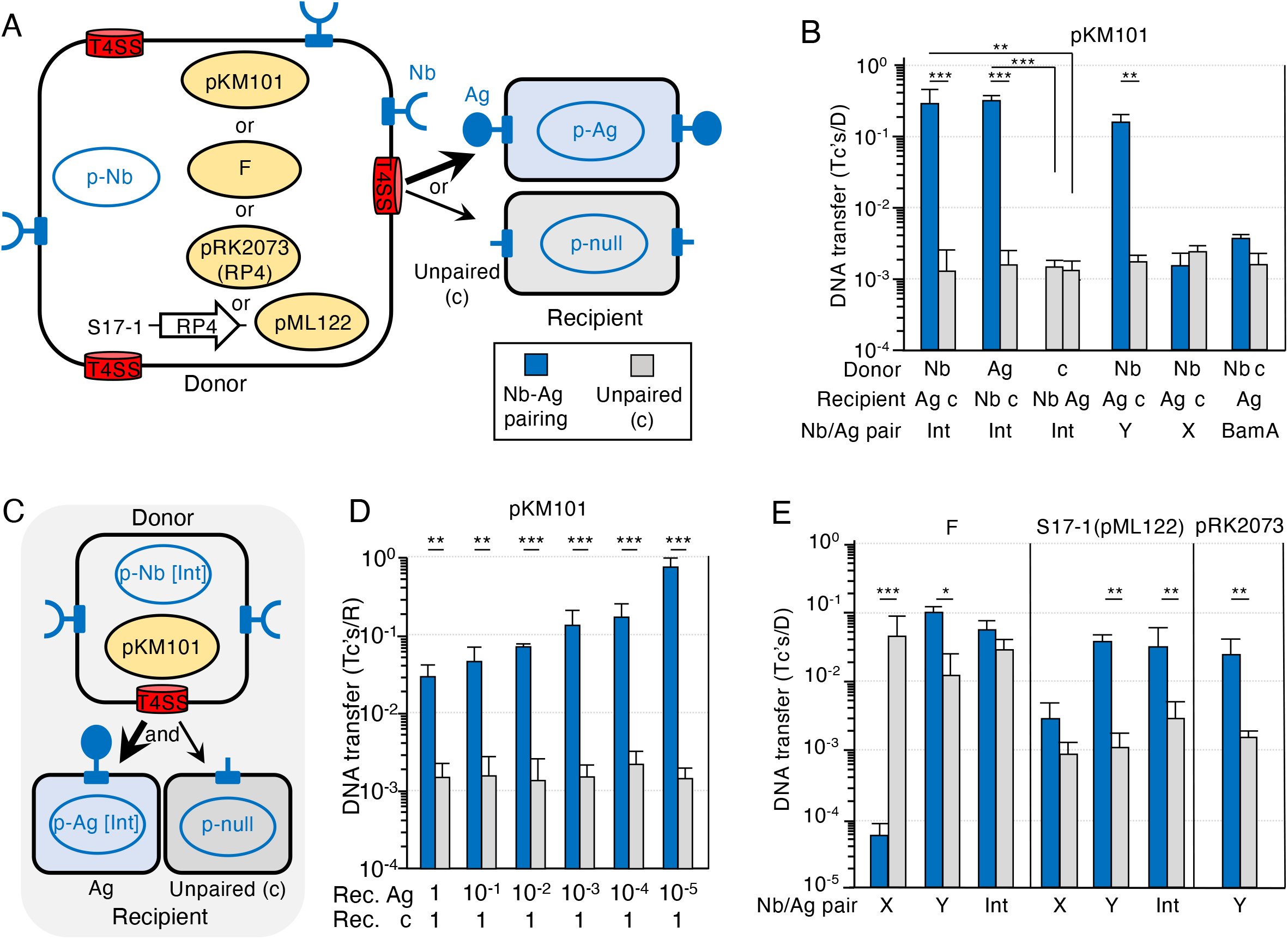
Nb/Ag pairing promotes targeted plasmid transfer. **A.** Schematic depicting *E. coli* MC4100 or S17-1 donor cells with surface-displayed Nb’s produced by p-Nb plasmids and T4SSs elaborated by one of the conjugation systems shown. MC4100-Chl recipients display cognate Ag’s or null control (c) produced by the respective p-Ag or p-null plasmids. Plasmids delivered to recipient cells are shaded in yellow. **B.** pKM101 transfer frequencies presented as transconjugants/donor (Tc’s/D). pKM101-carrying donors producing an Nb or Ag, or c (null control) were mated with recipients producing the cognate Ag or Nb, or c (null control). Specific Nb/Ag pairs (X, Y, Int, BamA) are listed at the bottom. Mating conditions: 60 min liquid, 1::1 D::R seed ratio, ATc 10^2^ ng/ml (final conc.). **C.** Schematic depicting mixed matings. **D.** MC4100 donors producing Nb [Int] from the p-Nb plasmid and harboring pKM101 were mixed with two recipient strains, MC4100-Rif producing Ag [Int] (Ag) or MC4100-Chl producing the null control (unpaired) at the seed ratios shown at the bottom (donor seed ratio of 1); other mating conditions as described in panel **B**. Plasmid transfer frequencies presented as Tc’s/R to allow for comparisons between the two recipient strains. **E.** Transfer frequencies of the F plasmid (pOX38), S17-1 mobilization of IncQ plasmid pML122 through the chromosomally-encoded RP4 T4SS, and pRK2073 self-transfer through the RP4 T4SS. Donors producing the Nb shown were mated with recipients producing the cognate Ag or c (null control). Mating conditions: 30 min liquid, 1::1 D::R seed ratio, ATc 10^2^ ng/ml (final conc.). All matings were repeated at least three times in triplicate, and the average transfer frequencies were presented as blue or gray bars with standard deviations shown as error bars. *p*-values between indicated data sets were calculated by the homoscedastic Student’s *t*-test. * p<0.05, ** p<0.001, *** p<0.0001.

We initially observed that Nb/Ag [Int] pairing conferred appreciably more aggregation of pKM101-carrying donor and recipient cells than the other three pairs (**Fig. S1A**). In the absence of Nb/Ag pairing, *E. coli* donors transferred pKM101 at frequencies of ∼10^3^ transconjugants/donor (Tc’s/D) in 60 min liquid matings (**Fig. S1B**). Strikingly, upon Nb/Ag [Int] pairing, donors delivered pKM101 to recipients at ∼10^2.5^-fold higher frequencies than to unpaired recipients (**Fig. 1B, S1B**). Nb/Ag [Int] pairing enhanced pKM101 transfer regardless of whether donors displayed the Nb or Ag and recipients the cognate construct (**Fig. 1B**). Nb/Ag [Y] pairing also induced appreciable cellular aggregation of pKM101 donor and paired recipients (**Fig. S1A**) and, correspondingly, enhanced pKM101 transfer by >10^2^-fold. However, neither Nb/Ag [X] nor Nb/Ag [BamA] induced appreciable aggregation of pKM101 donors (**Fig. S1A**), and neither conferred detectable stimulatory effects on pKM101 transfer (**Fig. 1B**).

We next examined the efficiency of Nb/Ag pairing on plasmid transfer in mixtures of Nb [Int] donors and both Ag [Int] and null recipient cells (**Fig. 1C**). Nb [Int] donors and null recipients were mixed at ratios of 1::1 colony-forming-units (CFUs), while Ag [Int] recipients were added at 10-fold serial dilutions of 1 to 10^-5^ CFUs. Nb [Int] donors transferred pKM101 to the null recipients at similar frequencies of ∼10^-3^ Tc’s/D regardless of Ag [Int] cell numbers (**Fig. 1D**). Remarkably, although ∼1 in 50 Ag [Int] recipient cells acquired pKM101 in matings with a 1::1::1 seed ratio, nearly every Ag [Int] recipient cell acquired the plasmid when present at 10^-5^ CFUs relative to Nb [Int] donor and null recipient cells. Nb/Ag pairing thus strongly enhances plasmid transfer to targeted recipients even when these recipients are present in very low numbers relative to untargeted cell populations.

The experimental conditions for assessing effects of Nb/Ag pairing on pKM101 transfer in the above experiments were derived from optimization of various parameters (**Fig. S1**). In time course studies, we observed that the magnitude of the stimulatory effect of Nb/Ag [Int] pairing on pKM101 transfer was greatest (∼10^2.5^-fold) in 60 min liquid matings, although interestingly pairing enhanced pKM101 transfer by nearly 10^1^-fold in matings as brief as 5 min and also significantly enhanced transfer in longer matings of 270 min even though background levels of transfer to the null recipient had also increased (**Fig. S1B**). In solid-surface matings, Nb/Ag pairing did not stimulate pKM101 transfer beyond the high frequencies (∼10^0^ Tc’s/D) observed in the absence of pairing; however, we did detect a nearly 10^1^-fold stimulatory effect of pairing in brief 5 min matings (**Fig. S1B**). We also varied the donor (D) to recipient (R) input ratio as determined by CFU counts, and determined that Nb/Ag [Int] pairing elicited the greatest stimulatory effect on pKM101 transfer with a D::R ratio of 1::1 (**Fig. S1C**). Finally, we varied the concentration of anhydrotetracycline (ATc), the inducer for expression of the Nb, Ag, or null constructs, and found that ATc at 10^2^ ng/ml (final conc.) yielded the greatest stimulatory effects on pKM101 transfer (**Fig. S1D**). The highest tested dose of ATc (10^3^ ng/ml) decreased DNA transfer, possibly due to toxicity resulting from overproduction of the Nb or Ag constructs. Based on these findings, we herewith assessed effects of Nb/Ag pairing on pKM101 transfer with 60 min liquid matings, a D::R seed ratio of 1::1, and ATc inducer at 10^2^ ng/ml (final conc.).

### Nb/Ag pairing stimulates F- and RP4-mediated transfer

Next, we tested whether Nb/Ag pairing stimulates transfer through other conjugation systems, including those encoded by the classical F plasmid (IncF) and RP4 (IncP) (**Fig. 1A**). To monitor F transfer, we used pOX38::Tc (Table S1), a widely studied derivative of F bearing a *tetR* gene (hereafter termed F). In contrast to pKM101 host cells, which elaborate brittle N pili that either break or are actively sloughed from the cell surface, F-carrying cells elaborate long and flexible retractile F pili that facilitate high frequency transfer even in aqueous environments (20, 21). In liquid matings of 30 min, *E. coli* donors transferred the F plasmid at ∼10^-2^ Tc’s/D in the absence of Nb/Ag pairing (**Fig. 1E**). Of the tested Nb/Ag pairs, Nb/Ag [Y] elicited the greatest aggregation in mixes of paired F-carrying donor and recipient cells (**Fig. S2A**). Correspondingly, Nb/Ag [Y] pairing stimulated F transfer by nearly 10^1^-fold in 30 min liquid matings (**Fig. 1E**). Stimulatory effects were even greater in brief 5 min liquid matings, although results of experimental repetitions were more variable, and negligible with matings of 60 or 270 min **(Fig. S2B)**. In 60 min solid-surface matings favoring high-frequency F transfer, Nb/Ag pairing did not detectably elevate transfer beyond frequencies achieved without pairing (**Fig. S2B**). We also evaluated effects of other Nb/Ag pairs, and found that Nb/Ag [Int] pairing did not significantly stimulate F transfer, whereas strikingly Nb/Ag [X] pairing conferred a strong inhibitory effect as evidenced by a ∼10^3^-fold reduction in transfer in 30 min liquid matings **(Fig. 1E).** Hereafter, we thus deployed the Nb/Ag [Y] pair and 30 min liquid matings (1::1 D::R seed ratio, 10^2^ ng/ml ATc) to assess effects of Nb/Ag pairing on F transfer.

We deployed two *E. coli* donor strains, S17-1 (22) and MC4100(pRK2073) (23), to assess effects of Nb/Ag pairing on RP4-mediated transfer. Strain S17-1 carries RP4 integrated in its chromosome and is widely used as a helper strain (22) to mobilize the transfer of nonself-transmissible plasmids, e.g., IncQ RSF1010 or its derivative pML122 (Table S1). pRK2073 is a ColE1 plasmid with cloned RP4 Tra and *oriT* regions, and therefore is self-transmissible (23). With these two donor strains, we were therefore able to directly compare the effects of Nb/Ag pairing on RP4-mediated mobilization vs self-transfer. As observed with the F system, Nb/Ag [Y] pairing induced appreciable aggregation of S17-1(pML122) and MC4100(pRK2073) donors (**Fig. S2C**). Interestingly, Nb/Ag [Y] pairing strongly enhanced pML122 mobilization in liquid matings of all tested durations (5 - 270 min), with the 30 min matings yielding the greatest ∼10^1.5^-fold stimulatory effect (**Fig. 1E, S2D**). Nb/Ag [Y] pairing did not detectably stimulate pML122 transfer in solid-surface matings (**Fig. S2D**). Among the three Nb/Ag pairs tested, Nb/Ag [Y] and Nb/Ag [Int] strongly enhanced pML122 transfer while Nb/Ag [X] had no significant effect (**Fig. 1E**). We used mating conditions optimized for S17-1 mobilization of pML122 (Nb/Ag [Y] pairing, 30 min liquid matings, 1::1 D::R seed ratio, ATc 10^2^ ng/ml) to assess effects of Nb/Ag pairing on pRK2073 self-transfer (**Fig. 1E).** Interestingly, MC4100(pRK2073) donors transferred pRK2073 at frequencies comparable to S17-1-directed transfer of pML122 and, moreover, Nb/Ag [Y] pairing enhanced pRK2073 transfer by ∼10^1.5^-fold, similar to the effect on pML122 mobilization (**Fig. 1E**).

### Nb/Ag pairing substitutes for T4SS-associated surface adhesins, but not conjugative pili

Conjugative elements encode pili or other surface adhesins that play important roles in establishment of target cell contacts (24). We tested whether Nb/Ag [Int] pairing could functionally replace the pKM101-encoded N pilus, but pKM101Δ*traM* donors lacking the TraM pilin subunit (18) were completely defective for plasmid transfer in liquid or solid surface matings regardless of Nb/Ag [Int] pairing (**Fig. 2A, B**). Efficient pKM101 transfer additionally requires two adhesins, TraC and Pep (**Fig. 2A**) (19). TraC is required for elaboration of N pili, likely localizing at the pilus tip where it promotes attachment to target cells (17, 18, 25). Alternatively, TraC is exported to the donor cell surface, where it forms higher-order adhesin complexes independently of and together with another pKM101-encoded protein termed Pep (19). Nb/Ag [Int] pairing did not restore transfer proficiency of donor cells harboring a Δ*traC* mutation in pCGR125, a plasmid that carries the entire Tra region of pKM101 (Table S1), either in liquid or solid-surface matings (**Fig. 2B**). Strikingly, however, Nb/Ag [Int] pairing strongly enhanced by >10^4^-fold the conjugation proficiency of Δ*pep* mutant donors, yielding mating frequencies even greater than observed for transfer of WT pKM101 (**Fig. 2B**). The Pep adhesin also contributes a ∼10^2^-fold enhancement in transfer of pKM101 in solid-surface matings (Fig. 2B) (19), and under these mating conditions Nb/Ag [Int] pairing also significantly enhanced the conjugation proficiency of Δ*pep* mutant donors (**Fig. 2B**). Nb/Ag [Int] pairing thus fully substitutes for the Pep surface adhesin, presumably by enhancing formation of pKM101 mating junctions.

**Fig. 2.**
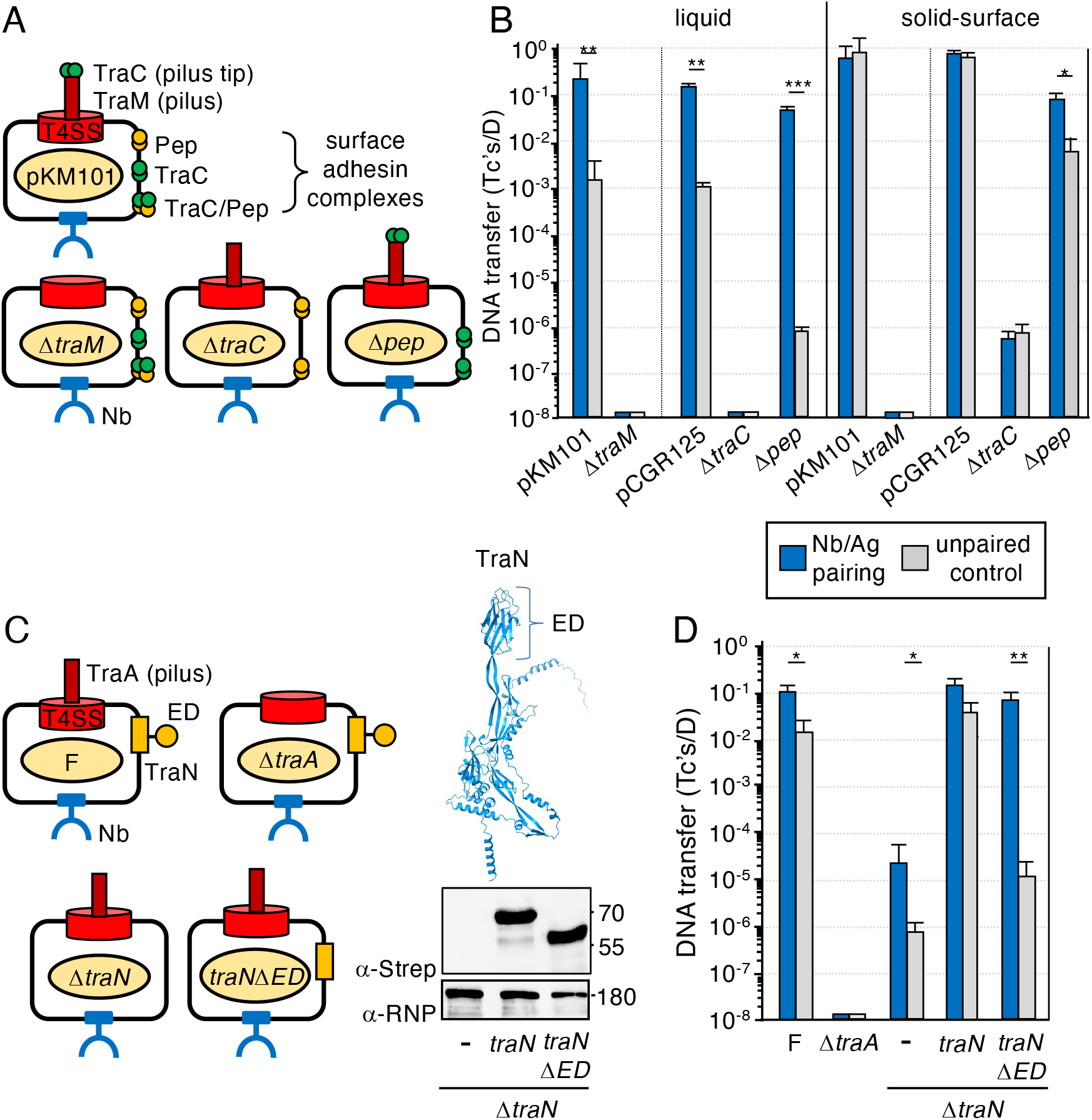
Nb/Ag pairing substitutes for T4SS-associated surface adhesins but not conjugative pili. **A.** Schematic of MC4100 donors harboring pKM101 or variant plasmids deleted of the genes shown. Strains additionally carry a p-Nb plasmid (not shown) for surface display of Nb [Int]. **B.** Frequencies of transfer through the pKM101 T4SS by MC4100 donors shown in panel **A**. MC4100-Rif recipients produced Ag [Int] or null control. pCGR125 is a pBAD24 plasmid carrying a functional *tra* region of pKM101, and pCGR125 variants are deleted of *pep* or *traC*; these mutations are fully complemented by expression *in trans* of the corresponding genes (19). Functionality of pCGR125 donors was assessed by mobilization of the pKM101 *oriT* plasmid pJG142 (Table S1), which transfers at frequencies equivalent to pKM101 (23). Plasmid transfer frequencies presented as Tc’s/D. Mating conditions: 60 min liquid or solid-surface, seed ratio 1::1, ATc 10^2^ ng/ml. **C.** Left: Schematic of MC4100 donors harboring the F plasmid pOX38 or variant plasmids deleted of the genes or domains shown; the Δ*traA* and Δ*traN* mutations are fully complemented by *trans-*expression of the corresponding genes (27). Strains additionally carry a p-Nb plasmid (not shown) for surface display of Nb [Y]. Right: Structure of TraN as predicted by Alphafold (73) with the extracellular domain (ED) highlighted. Production of TraN-Strep and TraNΔED-Strep by MC4100(pOX38Δ*traN*) without or with pBAD plasmids expressing the *traN* variants (Table S1) was assessed by immunostaining of western blots with α-Strep antibodies. Immunodetection of the β-subunit of *E. coli* RNA polymerase (RNP) served as a loading control. Molecular sizes (in kDa) of protein markers are listed at the right. **D.** Frequencies of transfer through the F T4SS by MC4100 donors depicted in panel **C.** MC4100-Rif recipients produced Ag [Y] or null control. Mating conditions: 30 min liquid, seed ratio 1::1, ATc 10^2^ ng/ml). Panels **B & D:** Mating replicates and statistical analyses as described in Fig. 1.

F plasmids transfer at high frequencies in dilute or dense cell growth conditions through elaboration of retractile F pili (21) and by stabilization of mating pairs, which is achieved through binding of donor-encoded TraN to outer membrane protein (OMP) receptors carried by recipient cells (**Fig. 2C)** (26, 27). Reminiscent of our findings with the pKM101 system, Nb/Ag [Y] pairing did not restore transfer proficiency of F-carrying donors deleted of the F-encoded TraA pilin gene in liquid or solid surface matings **(Fig. 2C, D).** Nb/Ag [Y] pairing also only partially compensated for the absence of TraN **(Fig. 2D)**, which is not surprising given that TraN contributes in various ways to elaboration of retractile F pili and T4SS channel function (26, 27). However, TraN’s role in mating pair stabilization is conferred by an extracellular domain (ED) that mediates the OMP interaction (**Fig. 2C**) (28, 29). We thus deleted TraN’s ED, first showing that Strep-tagged forms of TraNΔED mutant and TraN produced from isogenic plasmids accumulated at similar levels **(Fig. 2C, right).** Deletion of TraN’s ED and, by extension, the TraN - OMP interaction, attenuated F transfer by ∼10^3^-fold. Remarkably, upon Nb/Ag [Y] pairing, TraNΔED mutant donors regained transfer proficiency to levels observed for the WT F plasmid (**Fig. 2D**). These findings show that Nb/Ag [Y] pairing fully compensates for the function of TraN specifically related to stabilization of mating junctions.

### Nb/Ag pairing enhances targeted delivery and killing by pCrispr plasmids

Several studies have recently shown that conjugative plasmids equipped with CRISPR/Cas9 systems, which induce double-stranded breaks (DSB’s) in target sequences (30), can be deployed to cure antibiotic resistance plasmids, modify genomes, and kill targeted bacterial species (30–36). To determine whether Nb/Ag pairing enhances conjugation-mediated killing of target cells bearing CRISPR/Cas9 target sequences, we constructed two pCrispr plasmids. One, designated pCrispr*_chlR_*, carries a CRISPR/Cas9 element equipped with two gRNA sequences specific for distinct targets within the *chl^R^*gene in *E. coli* strain MC4100-Chl. The second, pCrispr_gRNA-_, harbors the CRISPR/Cas9 element bearing a non-specific gRNA sequence (negative control). We then introduced *oriT* sequences from the pKM101, F, or RP4 plasmids into both pCrispr plasmids for mobilization by the respective conjugation systems (**Fig. 3A**).

**Fig. 3.**
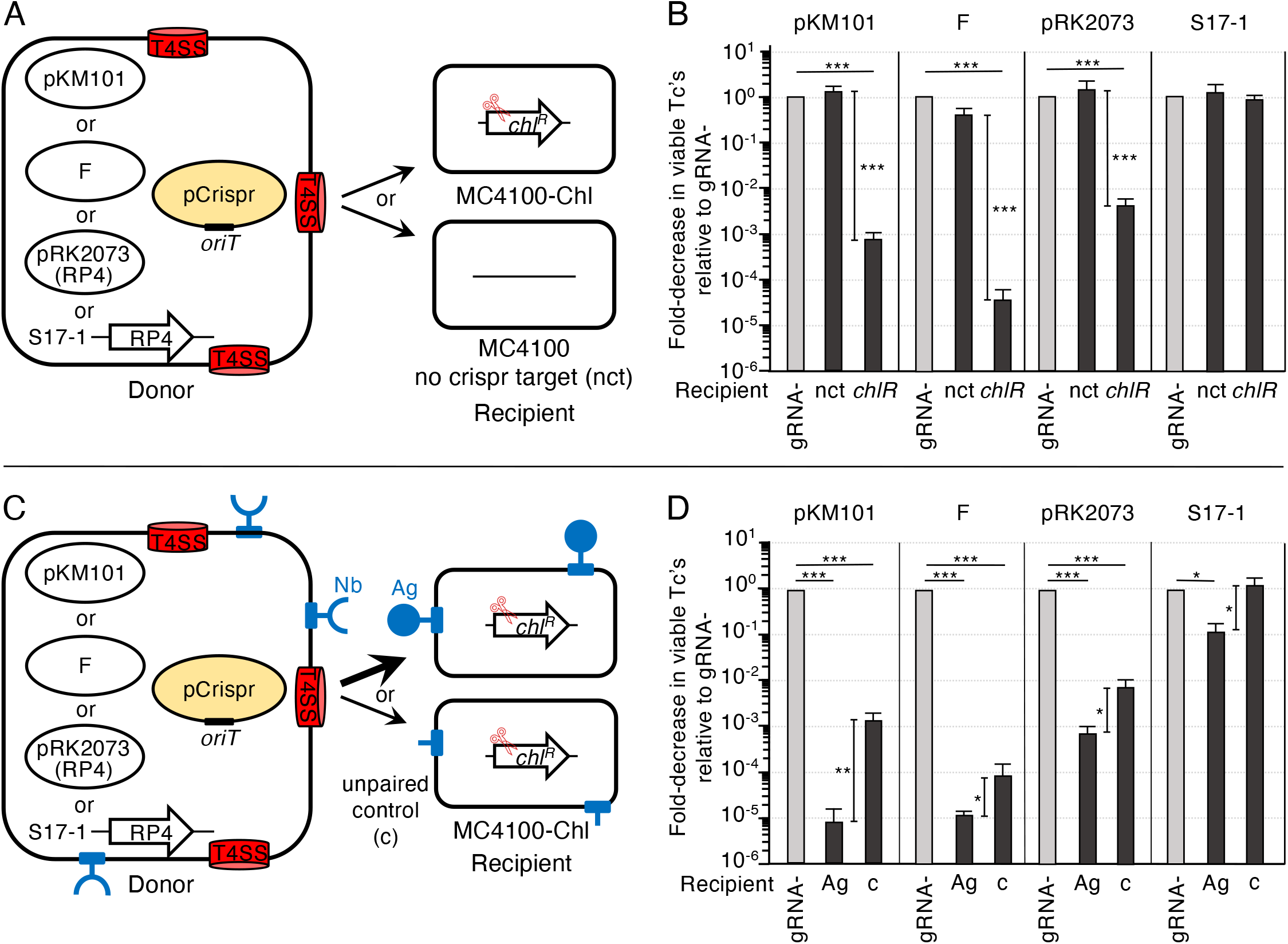
Effects of Nb/Ag pairing on pCrispr plasmid transfer and killing of *E. coli* recipients. A. Schematic depicting *E. coli* MC4100 or S17-1 donor cells with T4SSs elaborated by one of the conjugation systems shown, and pCrispr*_chlR_* (targets *chl^R^* gene) or pCrispr_gRNA-_ (nonspecific guide RNA control) harboring cognate *oriT* sequences (plasmids represented by pCrispr shaded in yellow). MC4100-Chl recipients harbor the CRISPR/Cas9 target sequence (*chl^R^*, scissors), MC4100 recipients lacking the *chl^R^*target gene are denoted nct (<u>n</u>o <u>c</u>rispr target). Both recipients are Rif^R^. **B.** Killing efficiencies presented as the fold-decrease in viable transconjugants (Tc’s) resulting from transfer of pCrispr*_chlR_* relative to viable Tc’s resulting from transfer of pCrispr_gRNA-_ (normalized to 1 and represented by gray bars for each of the conjugation systems). Matings with pKM101 donors were carried out for 60 min and for the F and RP4 donors for 30 min in liquid (1::1 D::R seed ratio, ATc 10^2^ ng/ml). Matings were repeated at least three times in triplicate with average killing efficiencies presented as black bars. Standard deviations shown as error bars. Statistical significance values at top represent comparisons of viable Tc’s resulting from transfer of the pCrispr_chlR_ plasmids relative to gRNA-plasmids (normalized to 1). Values presented vertically represent comparisons of viable Tc’s resulting from transfer of the pCrispr*_chlR_*plasmids into the nct and *chlR* recipient strains. * p<0.05, ** p<0.001, *** p<0.0001. **C.** Schematic as in Panel A, but with MC4100 donors additionally producing Nb’s. Donors were mated with MC4100-Chl recipients producing the cognate Ag’s or the null control. **D.** Results of matings between Nb-producing donors and recipients producing the cognate Ag or null control. Nb/Ag pairing: [Int] for pKM101-mediated matings; [Y] for F- and RP4-mediated matings. Other mating conditions, killing efficiencies and statistical analyses were as described in Panel **B**.

In liquid matings, all three conjugation systems mobilized the transfer of pCrispr*_chlR_* and pCrispr_gRNA-_plasmids at equivalent frequencies to MC4100 recipients (designated as the <u>n</u>o <u>c</u>rispr target or nct strain) (**Fig. 3B**). By contrast, delivery of pCrispr*_chlR_* to isogenic MC4100-Chl bearing the *chlR* target sequence (designated *chlR* strain) elicited significant levels of killing of transconjugants relative to transfer of the pCrispr_gRNA-_plasmids (**Fig. 3B**). For pKM101-carrying donors, pCrispr*_chlR_* mobilization elicited a ∼10^3^-fold reduction in viable MC4100-Chl transconjugants compared with transfer of the gRNA-control plasmid. F-carrying donors elicited a striking >10^4^-fold killing effect, whereas pRK2073-carrying donors elicited a ∼10^2^-fold killing effect. S17-1 donors with chromosomally-integrated RP4 delivered both pCrispr plasmids to MC4100 recipients at equivalent frequencies, but curiously the transfer of pCrispr*_chlR_*to MC4100-Chl recipients did not induce killing (**Fig. 3B**).

For each of the tested conjugation systems, including the S17-1-encoded RP4 system, Nb/Ag-pairing significantly enhanced CRISPR/Cas9-mediated killing of paired vs unpaired (null) MC4100-Chl transconjugants (**Fig. 3C**). Whereas pKM101-mediated transfer of pCrispr*_chlR_* elicited a ∼10^3^-fold killing of MC4100-Chl recipients in the absence of Nb/Ag pairing, Nb/Ag [Int] pairing conferred an additional >10^2^-fold killing effect (**Fig. 3D**). This increase in killing is in line with the increment of the stimulatory effect observed for Nb/Ag [Int] pairing on pKM101 transfer (**see Fig. 1B**). For the F and both RP4 systems, Nb/Ag [Y] pairing similarly enhanced killing by 10^1^-fold, which also is in line with the observed stimulatory effects of Nb/Ag pairing on transfer of F, pRK2073, or S17-1-directed mobilization of pML122 (**see Fig. 1E**). Together, these findings establish that Nb/Ag pairing significantly enhances conjugation-mediated killing of cells harboring CRISPR/Cas9 target sequences over killing effects achieved in the absence of pairing.

### Minicells conjugatively transfer plasmids and kill recipient cells, but Nb/Ag-enhanced binding does not stimulate minicell-mediated transfer or killing

With a goal of further empowering our PDS, we asked whether *E. coli* minicells can kill intact cells through conjugative delivery of pCrispr plasmids. *E. coli* strains with defective Min systems generate anucleate minicells through mislocalization of division sites near the cell poles (37). Minicells retain many functions of intact cells, but are non-viable and therefore safer than intact cells for use as delivery vectors in infection settings. We first tested whether Nb/Ag pairing promotes binding of minicells purified from the *minD* mutant WM3886 (38) to intact *E. coli* cells (**Fig. 4A, left**). Remarkably, minicells with surface-displayed Nb [Int] abundantly bound *E. coli* MC4100 cells displaying Ag [Int], but only sporadically bound null cells (**Fig. 4A, middle**). The Nb [Int] minicells characteristically distributed in many copies around Ag [Int] cell peripheries. These minicells also induced appreciable aggregation of Ag [Int] cells but not of null cells, suggesting that they display multiple copies of Nb [Int] capable of binding Ag [Int] displayed by different cells. Among 100 Ag [Int] cells examined, an average of 5 Nb [Int] minicells bound per cell, although a few cells had ∼20 - 30 bound minicells. Among 100 null cells examined, an average of 1 minicells bound per cell, and the upper range of bound minicells was only 3 - 4 (**Fig. 4A, right**).

**Fig. 4.**
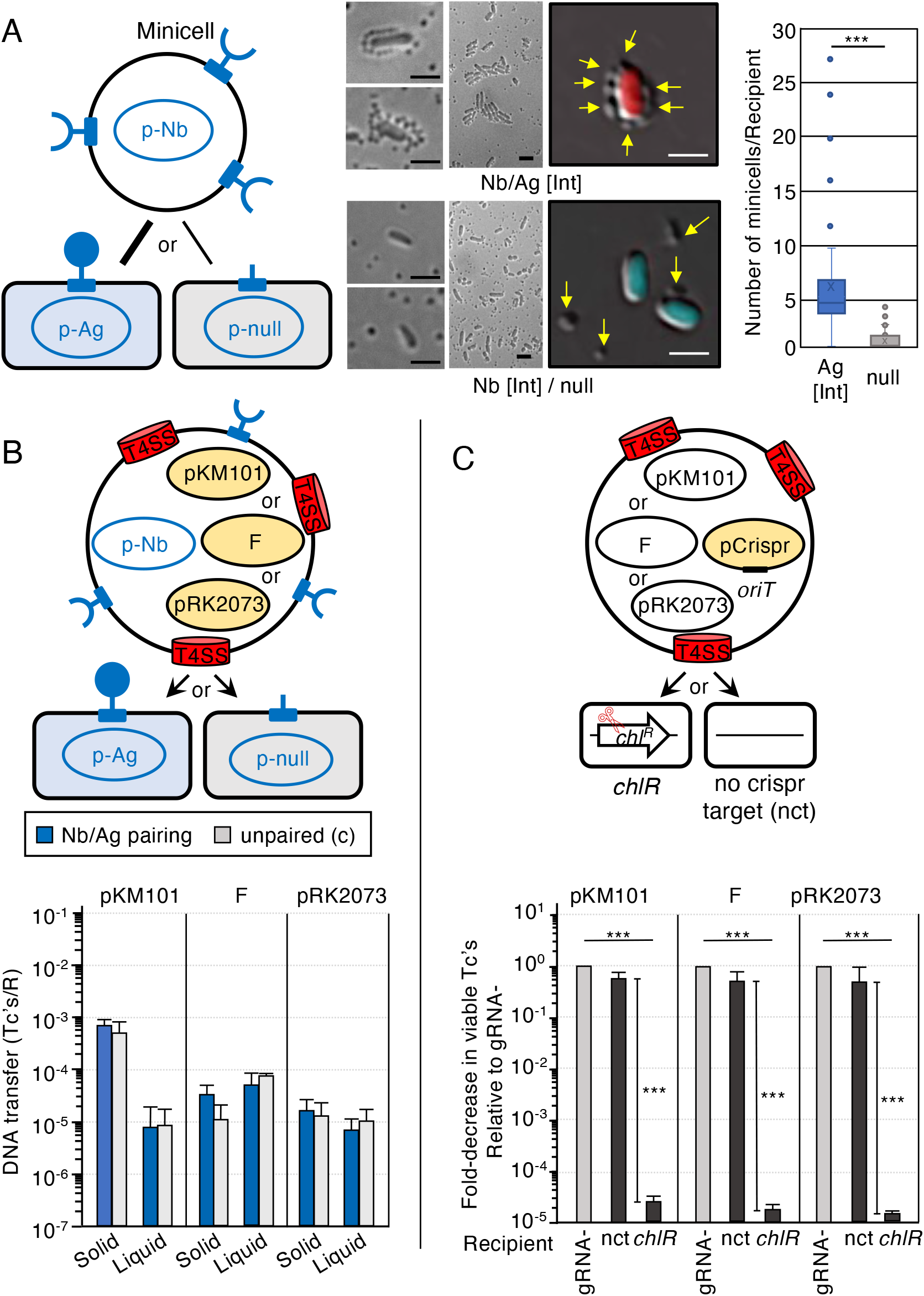
Nb/Ag pairing enhances minicell - cell binding, but not DNA transfer or killing. **A. Left:** Schematic of *E. coli* minicells purified from WM3886 producing Nb [Int] mixed with intact *E. coli* MC4100 cells producing Ag [Int] or the null control. **Middle:** Representative DIC images of Nb/Ag-paired vs unpaired mixtures. Intact *E. coli* cells produced mCherry (upper) or mCerulean3 (lower) for enhanced detection. Arrows point to minicells. Scale bars, 2 μm. **Right**: Box and Whisker plot showing numbers of Nb [Int] minicells bound to Ag [Int] or null *E. coli* cells (n=100 for both cell types). Box delineates upper and lower quartiles with X as the mean and line as the median of whole; whiskers denote maximum and minimum values with outliers shown as dots. **B. Upper:** Schematic depicting *E. coli* minicells with surface-displayed Nb’s produced by p-Nb and T4SSs elaborated by one of the conjugation systems shown. MC4100-Chl recipients display cognate Ag’s or null control produced by the respective p-Ag or p-null plasmids. Plasmids delivered to recipient cells are shaded in yellow. **Lower:** Plasmid transfer frequencies presented as transconjugants/recipient (Tc’s/R). Minicell donors with surface-displayed Nb’s were mated with MC4100-Chl recipients displaying cognate Ag’s or null control in liquid or on solid surfaces for 2 h. Nb/Ag pairing: [Int] for pKM101 transfer and [Y] for F and pRK2073(RP4) transfer; 10::1 minicell::recipient cell ratio. Mating replicates and statistical analyses as described in Fig. 1. No statistical differences between Nb/Ag-paired vs unpaired matings. **C. Upper:** Schematic depicting *E. coli* minicells with T4SSs elaborated by one of the conjugation systems shown, and pCrispr*_chlR_* or pCrispr_gRNA-_harboring cognate *oriT* sequences (represented by pCrispr; shaded in yellow). MC4100-Chl recipients harbor the CRISPR/Cas9 target sequence (*chlR*, scissors), MC4100 recipients lacking the *chlR* target gene are denoted nct (no crispr target). Both recipients are Rif^R^. **Lower:** Killing efficiencies presented as the fold-decrease in viable transconjugants (Tc’s) resulting from transfer of pCrispr*_chlR_* relative to viable Tc’s resulting from transfer of pCrispr_gRNA-_ (normalized to 1 and represented by gray bars). Matings, killing efficiency representations, and statistical analyses as described in Fig. 3B.

Minicells purified from strains harboring pKM101, F or pRK2073 were proficient for transfer of these plasmids to intact *E. coli* recipients in both solid-surface and liquid matings (**Fig. 4B**). Specifically, minicells equipped with the F and pRK2073 systems delivered these plasmids at comparable frequencies of ∼10^-4^ - 10^-5^ Tc’s/R in both mating conditions. Minicells equipped with pKM101 transferred the plasmid at ∼10^-5^ Tc’s/R in liquid matings, but at an appreciably higher frequency of ∼10^-3^ Tc’s/R in solid-surface matings. Surprisingly, minicells equipped with any of the conjugation systems and displaying Nb’s showed no enhancement in plasmid transfer to the cognate Ag recipient in either solid-surface or liquid matings (**Fig. 4B**). Thus, despite clear stimulatory effects of Nb/Ag pairing on minicell - cell binding (**Fig. 4A**), the Nb/Ag-mediated interactions did not translate to elevated conjugative transfer frequencies.

In spite of these findings, we tested for minicell-mediated killing of intact recipient cells (**Fig. 4C**). At the outset, we showed that minicells equipped with the pKM101-, F-, or pRK2073(RP4)-encoded conjugation systems mobilized the transfer of both pCrispr*_chlR_* or pCrispr_gRNA-_plasmids to the no crispr target (nct) strain MC4100 at comparable frequencies. Strikingly, minicell-mediated mobilization of pCrispr*_chlR_* plasmids through each of the three conjugation systems elicited a nearly 10^5^-fold reduction in viable MC4100-Chl transconjugants relative to pCrispr_gRNA-_transfer frequencies. Minicells armed with the pKM101, F, or RP4 conjugation machines are therefore highly effectively in pCrispr mobilization and selective killing of cells harboring CRISPR/Cas9 target sequences, even in the absence of Nb/Ag pairing.

### Nb/Ag pairing enhances plasmid transfer and killing of *Pseudomonas aeruginosa*

Finally, we tested whether the *E. coli*-based PDS can be deployed to kill another species, namely, the opportunistic pathogen *P. aeruginosa*. Strain PAO-1Δ*tssB1* served as a recipient because it is defective for production of the type 6 secretion system (T6SS), which has been shown to target and kill *E. coli* strains harboring T4SSs (18, 39). Prior work has shown that pKM101 transfers poorly relative to RP4, but both plasmids are stably maintained in *P. aeruginosa* (40–42), whereas F fails to transfer or is unstable in this species (43). In solid-surface matings, we found that all three conjugation systems were proficient for mobilization of pML122 to *P. aeruginosa* recipients (**Fig. 5A**). For F-mediated transfer, we expressed a variant of the TraD substrate receptor deleted of a C-terminal discrimination motif (*traDΔC15*) that was shown to promote F transfer while inhibiting mobilization of IncQ plasmids (44–46). Nevertheless, *E. coli* donors harboring either of the pKM101 and F systems mobilized pML122 at low frequencies of 10^-7^ - 10^-8^ Tc’s/D in 5 h solid-surface matings, although transfer frequencies were higher (10^-3^ - 10^-4^ Tc’s/D) in long-term 20 h matings (**Fig. S3A**). S17-1 donors equipped with the RP4 system were considerably more proficient in mobilization of pML122 to *P. aeruginosa*, as evidenced by transfer frequencies approaching 10^-2^ Tc’s/D in 1 h matings and 10^0^ Tc’s/D in 4 h matings (Fig. S3A). MC4100(pRK2073) donors also robustly transferred pML122 in 1, 2, and 4 h matings, although transfer frequencies were routinely 1 – 2 orders of magnitude lower than observed for S17-1-mediated transfer at each of these mating times (**Fig. S3A**). This slight reduction in pML122 transfer might be attributable to competition between pML122 and pRK2073 for docking with and transfer through pRK2073-encoded T4SSs, which is absent in S17-1 donors harboring only the mobilizable pML122 substrate.

**Fig. 5.**
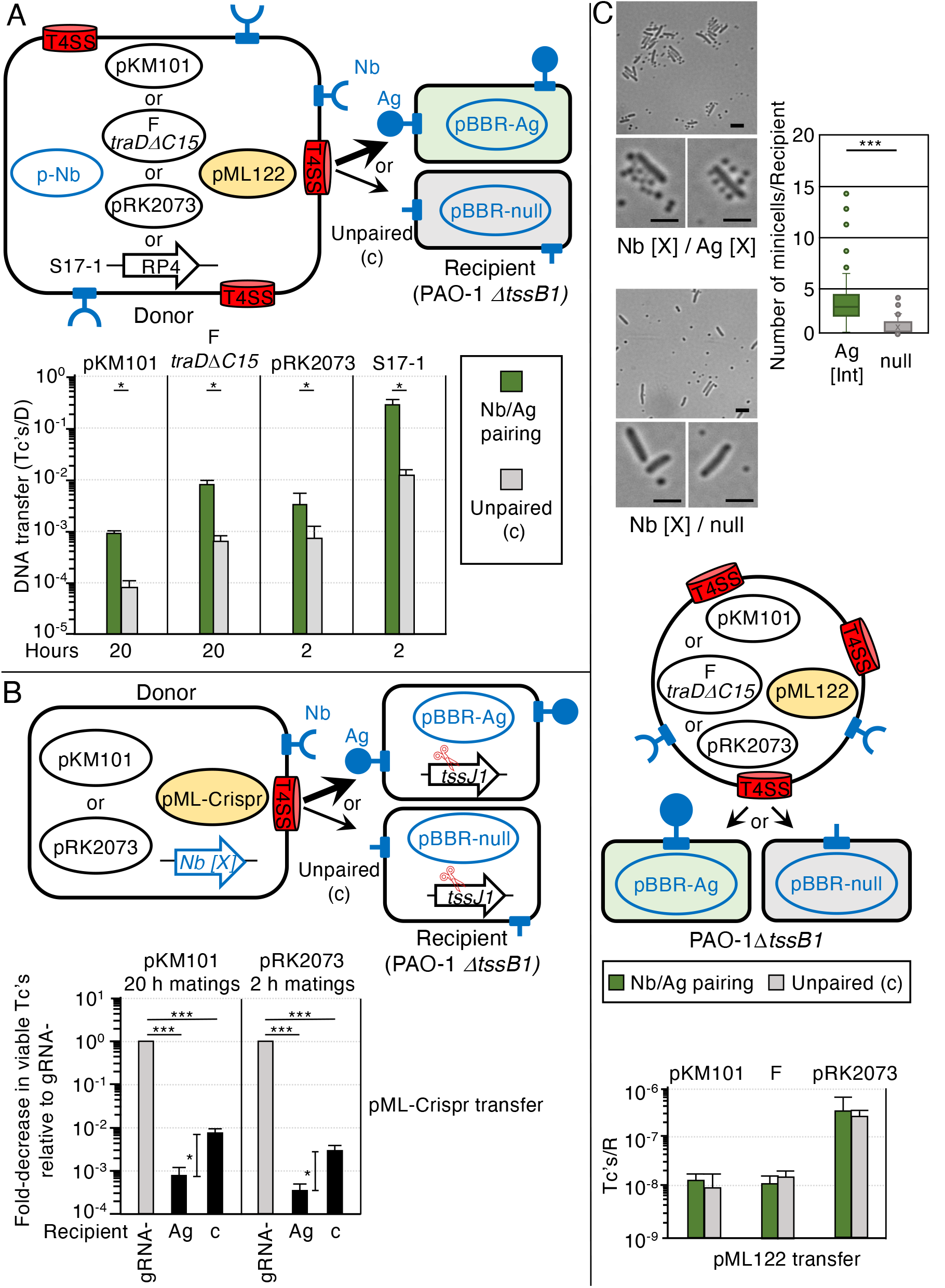
Nb/Ag pairing enhances *E. coli - P. aeruginosa* DNA transfer and killing. **A. Upper:** Schematic depicting *E. coli* MC4100 or S17-1 donor cells with surface-displayed Nb [X] produced by p-Nb and T4SSs elaborated by one of the conjugation systems shown. The F *traD*Δ*C15* oval represents pOX38Δ*traD* plus pYGL348, which expresses *traD*Δ*C15* from a salicylic acid-inducible promoter (Table S1). PAO-1Δ*tssB1* recipients display Ag [X] or null control produced by the respective pBBR-Ag or pBBR-null plasmids. Mobilizable IncQ pML122 is shaded in yellow. **Lower:** pML122 transfer frequencies in solid-surface matings of durations shown. Mating replicates and statistical analyses as described in Fig. 1. * p<0.05, ** p<0.001, *** p<0.0001. **B. Upper:** Schematic depicting *E. coli* MC4100 donor cells with surface-displayed Nb [X] produced by the chromosomally-integrated *Nb [X]* gene fusion, T4SSs elaborated by pKM101 or pRK2073, and mobilizable pCrispr*_tssJ1_* or pCrispr_gRNA-_ plasmids (represented by pML-Crispr; shaded in yellow). PAO-1Δ*tssB1* recipients harbor the CRISPR/Cas9 target sequence (*tssJ1*, scissors) and surface display Ag [X] or null control produced by the respective pBBR-Ag or pBBR-null plasmids. **Lower:** Killing efficiencies are presented as the fold-decrease in viable transconjugants (Tc’s) resulting from transfer of pCrispr*_tssJ1_* relative to viable Tc’s resulting from transfer of pCrispr_gRNA-_ normalized to 1 and represented by gray bars for each of the conjugation systems. Solid-surface matings of durations shown (1::1 D::R seed ratio, ATc 10^2^ ng/ml) were repeated at least three times in triplicate with average killing efficiencies presented as black bars. Standard deviations shown as error bars. Statistical significance as described in Fig. 3B. * p<0.05, ** p<0.001, *** p<0.0001. **C. Upper:** Representative DIC images of Nb/Ag-paired vs unpaired mixtures of *E. coli* minicells and intact *P. aeruginosa* cells. **Right**: Box and Whisker plot showing numbers of Nb [X] *E. coli* minicells bound to Ag [X] or null *P. aeruginosa* cells (n=100 for both cell types). Box delineates upper and lower quartiles with X as the mean and line as the median of whole; whiskers denote maximum and minimum values, outliers shown as dots. **Middle:** Schematic depicting *E. coli* minicells with surface-displayed Nb [X] produced by p-Nb (not shown) and T4SSs elaborated by one of the conjugation systems. F *traD*Δ*C15* oval is representative as described above. Mobilizable pML122 is shaded in yellow. PAO-1Δ*tssB1* recipients display Ag [X] or null control produced by the respective pBBR-Ag or pBBR-null plasmids. **Lower:** pML122 mobilization frequencies presented as transconjugants/recipient (Tc’s/R). Matings were carried out for 12 h on solid-surfaces, other mating conditions and statistical analyses as described in Fig. 4B. No statistical differences between paired vs unpaired matings.

We engineered *P. aeruginosa* PAO-1Δ*tssB1* strains to produce surface-displayed Ag or null proteins from pBBRMCS, which also is stably maintained in *Pseudomonas spp.* (47). Among the three tested Nb/Ag pairs, Nb/Ag [X] conferred the greatest stimulatory effects on pML122 transfer by all three conjugation systems (**Fig. S3B**), which was surprising given that Nb/Ag [X] pairing was noneffective or strongly inhibited the *E. coli* - *E. coli* matings (see **Fig. 1B, E**). In solid-surface matings of the durations specified, Nb/Ag [X] pairing elicited increases of ∼10^1^- to 10^1.5^-fold in pML122 transfer frequencies, regardless of which conjugation systems donors harbored for mobilization (**Fig. 5A**).

We next constructed two pML122 substrates equipped with CRISPR/Cas9 elements either bearing two distinct guide RNAs designed to target *tssJ1* in the T6SS locus (pML-Crispr*_tssJ1_*) or with a nonspecific guide RNA sequence as a control (pML-Crispr_gRNA-_). For unknown reasons, the pML-Crispr plasmids were toxic in *E. coli* donors that also carried the p-Nb plasmids. We therefore recombineered genes and upstream promoters for the surface-displayed Nb [X], Ag [X], or null proteins into the chromosomal *chl^R^* gene in strain MC4100-Chl. Initial studies confirmed that a mixture of *E. coli* strains producing cognate Nb/Ag [X] pairs aggregated more extensively than the Nb [X] and null strains (**Fig. S3C**). A mixture of *E. coli* Nb [X] donors and Ag [X]- producing *P. aeruginosa* PAO-1Δ*tssB1* recipients also aggregated, although not to the extent observed with the equivalent *E. coli - E. coli* pairing (**Fig. S3C**). *E. coli* carrying pOX38Δ*traD,* a p15a plasmid expressing *traD*Δ*C15*, chromosomally-encoded Nb [X], and either of the pML-Crispr plasmids still grew poorly, preventing a test for F-mediated killing of *P. aeruginosa.* However, the *E. coli* Nb [X] donor harboring pKM101 and pML-Crispr*_tssJ1_* showed no growth defects, and when mated with Ag [X]-paired *P. aeruginosa* recipients elicited ∼10^3^-fold killing of transconjugants compared with the transfer frequency of the nonkilling pML-Crispr_gRNA-_ **(Fig. 5B).** Without Nb/Ag [X] pairing, transfer of pML-Crispr*_tssJ1_* elicited a ∼10^2^-fold killing effect, showing that Nb/Ag [X] pairing stimulated killing by about 10^1^-fold. *E. coli* donors harboring pRK2073 and pML-Crispr*_tssJ1_*similarly elicited a ∼10^1^-fold enhancement of killing when Nb/Ag-paired with *P. aeruginosa* recipients vs. the unpaired matings. Reminiscent of our findings with *E. coli - E. coli* matings (**Fig. 3B**), therefore, Nb/Ag pairing of *E. coli* with *P. aeruginosa* elevated killing by increments similar to the stimulatory effects of pairing on pKM101- or pRK2073-directed mobilization of pML122.

We extended these investigations by testing whether *E. coli* minicells equipped with Nb’s selectively bind and deliver plasmid cargoes to *P. aeruginosa* cells. Minicells displaying Nb [X] readily bound to Ag [X]-paired but not unpaired *P. aeruginosa* cells. Among 100 Ag [X]-displaying *P. aeruginosa* cells examined, an average of 3.5 Nb [X] minicells bound per cell, and a few cells had ∼10 - 15 bound minicells. Among the 100 null cells examined, an average of 1 minicells bound per cell, and the upper range of bound minicells was only 3 - 4 (**Fig. 5C, right**). The Nb/Ag [X] pairing system was therefore highly effective in promoting *E. coli* minicell - *P. aeruginosa* interspecies interactions (**Fig. 5C, top**).

In matings with PAO-1Δ*tssB1* recipients (**Fig. 5C, middle**), minicell donors harboring each of the three conjugation systems mobilized pML122 transfer, albeit at low frequencies (10^-8^ to 10^-7^ Tc’s/R) **(Fig. 5C, bottom).** As shown for the intraspecies *E. coli* minicell - cell matings (**Fig. 4B**), Nb/Ag [X] pairing did not stimulate minicell-mediated transfer of pML122 to intact *P. aeruginosa* beyond levels observed with unpaired matings (**Fig. 5C, bottom**). We tested for minicell-mediated killing of *P. aeruginosa*, but minicells equipped with any of the conjugation systems failed to elicit detectable killing of *P. aeruginosa* transconjugants. Given the low transfer frequencies observed for minicell mobilization of pML122 into *P. aeruginosa*, we suspect that minicell-mediated killing, if it occurs, is below the threshold of detection.

## DISCUSSION

Toward development of a bacterial PDS, our initial studies underscored the importance of systematically optimizing parameters such as specific Nb/Ag pairs, mating times, donor::recipient ratios, and Nb/Ag production levels for each conjugation system and targeted species of interest. Most interestingly, we observed a range of effects of the different Nb/Ag pairs on conjugation, from strong enhancement of pKM101 and RP4 transfer to *E. coli* recipients in liquid matings to milder enhancement of plasmid mobilization to *P. aeruginosa* recipients in solid-surface matings, to strong inhibitory effects on F plasmid transfer to *E. coli* recipients. Although the goal of this work was to test the stimulatory effects of Nb/Ag pairing for conjugation-mediated intercellular killing, the finding that pairing can specifically inhibit given conjugation machines is also of practical value for efforts to thwart dissemination of clinically-relevant MGEs such as F plasmids and their cargoes of antibiotic resistance genes. We suspect that a combination of factors including Nb/Ag binding affinities, structural features of the Nb/Ag complex, and abundance of Nb/Ag pairs at the cell-cell interface can influence the impact of Nb/Ag pairing on conjugative transfer. Individual characteristics of conjugation systems, e.g., spatial localization, and features of the donor and target cell envelopes also potentially affect this outcome. With experimental parameters optimized for the three model conjugation machines, we demonstrated that specific Nb/Ag pairs significantly stimulated plasmid transfer through the pKM101 and RP4 T4SSs in liquid matings to the extent that transfer frequencies were elevated to levels achieved in solid-surface matings. We also showed that Nb/Ag pairing promotes robust pKM101 transfer even in matings in which targeted recipient cells are present at very low ratios compared with off-target recipients, e.g., 1::10^5^, and that under such conditions nearly every targeted recipient acquires the plasmid. In an infection setting, Nb/Ag pairing thus can be expected to significantly elevate targeted transfer even to a pathogenic species of interest that is present in very low numbers.

Overall, results of our initial studies agree with those of Robledo *et al.,* who presented the first evidence for stimulatory effects of Nb/Ag pairing on conjugative transfer of IncN, IncW, and IncP plasmids in liquid matings (48). Both studies also showed that Nb/Ag pairing is less effective in elevating transfer of F plasmids in liquid matings, presumably because these systems elaborate dynamic F pili that enhance the probability of mating pair formation in dilute cellular environments. Importantly, however, we identified several features or modifications of the F system of potential value for its widespread deployment in a PDS. First, we found that Nb/Ag pairing in fact confers a statistically-significant enhancement in F plasmid transfer when matings are carried out for brief times. Translating such a finding to a therapeutic setting, it is enticing to speculate that F machines could be rendered tunable for bursts of targeted transfer, for example, through tight regulation of the VirD4-like substrate receptor TraD. This could have the dual advantage of rapidly killing targeted cells while also minimizing inactivation of the delivered killing agent, e.g., a CRISPR/Cas9 system, through mutation (see below).

Second, by deletion of TraN’s ED, we essentially converted the F system into a poorly-functioning machine whose transfer proficiency could be fully restored by Nb/Ag pairing. In the context of Nb/Ag pairing, therefore, donors harboring the *traN*Δ*ED* mutation exhibit a low off-target transfer efficiency. This finding, coupled with our evidence that Nb/Ag pairing also elevated transfer by pKM101Δ*pep* mutant machines by ∼10^5^-fold, supports a general model that T4SSs rendered functionally impaired through mutation, whose robust activities are restorable by Nb/Ag pairing, are ideal for deployment in a PDS because they fulfill the two desired outcomes of a PDS - high-frequency targeted and low-frequency off-target transfer.

Third, while recent studies have shown that the TraN - OMP interaction plays a role in specifying the host range of given F plasmids among Enterobacterial species (29, 49), our finding that Nb/Ag pairing substituted for the TraN ED - OMP interaction in principle enables rewiring of the F system for conjugation with any cell type of interest. Supporting this idea, Nb/Ag [X] pairing significantly enhanced F-mediated pML122 mobilization from *E. coli* donors to *P. aeruginosa* recipients (**Fig. 5**). That Nb/Ag [X] pairing did not elevate interspecies mobilization of pML122 to a greater extent than ∼1 log could be due to relative inefficiency of the tested Nb/Ag pairs on aggregation of *E. coli* and *P. aeruginosa* cells as compared to *E. coli - E. coli* cells (**Fig. S3C**). The notion that Nb/Ag pairing empowers the rewiring of T4SSs gains additional support from recent evidence that the *A. tumefaciens* VirB/VirD4 T4SS, which in natural settings delivers oncogenic T-DNA to plant cells, can be repurposed through Nb/Ag pairing to deliver protein cargoes to yeast and mammalian cells (50).

With the fully assembled PDS, we showed for the first time that Nb/Ag pairing stimulates conjugative transfer between *E. coli* and *P. aeruginosa* species, and also significantly enhances conjugation-directed CRISPR/Cas9- mediated killing of *E. coli* and *P. aeruginosa* targets. Our findings strengthen prior claims that conjugative delivery of CRISPR/Cas9 systems constitutes an effective antimicrobial strategy (30–36, 51). Among the three model conjugation systems analyzed, we determined that the pKM101 system proved most effective for Nb/Ag-targeted killing of *E. coli* recipients, while the RP4 system elicited the greatest Nb/Ag-targeted killing of *P. aeruginosa* recipients. The latter finding is consistent with evidence that the RP4-encoded T4SS is highly versatile in its capacity to transfer DNA cargoes to most if not all Gram-negative bacterial species as well as fungi and mammalian cells (52–54). Specific features of the RP4-encoded T4SSs that remain to be identified might be especially well adapted for deployment in PDSs targeting phylogenetically diverse cell types.

Although we found that Nb/Ag pairing stimulates killing efficiencies by as much as 10^5^-fold, an appreciable number of transconjugants, e.g., ∼10^3^ among a total of 10^8^ transconjugants for the pKM101 system (**Fig. 3B**), escaped CRISPR/Cas9-mediated killing. Previous studies have attributed such ‘escape’ frequencies to mutational inactivation of the CRISPR/Cas9 elements or of target sequences, or repair of CRISPR/Cas-directed DSB’s (35). Natural mutation rates and host-encoded repair functions can account for these ‘escapes’, but it is also noteworthy that the act of conjugation itself potentially mitigates the killing effects accompanying CRISPR/Cas9 transfer. This is partly because MGEs have evolved various strategies to evade CRISPR systems, including production of anti-CRISPR proteins or other DNA-restriction, modification, or repair enzymes (55–60). Additionally, mating activates the SOS response and associated error-prone replication and DNA repair and recombination pathways through sensing of incoming ssDNA intermediates by new transconjugants (61–64). Indeed, this latter feature of conjugation might account for our finding that S17-1-mediated transfer of pCrispr*_chlR_*failed to elicit strong killing of *E. coli* recipients, since S17-1 is known to translocate its chromosome through processing at the integrated RP4 *oriT* (65), and such Hfr transfer is a known strong inducer of the SOS response (61). It is also important to point out that, in contrast to S17-1 donors which deliver non-self-transmissible plasmids to target cells, donors harboring the pKM101, F, or pRK2073 plasmids potentially deliver both the conjugative plasmid and pCrispr*_chlR_* plasmid to recipients, setting the stage for redundant transmission of both plasmids to new recipients and reiterative killing of recipients. In view of these observations, we suggest several possible improvements to our current PDS strategy. First, pCrispr plasmids deployed in a PDS should be deleted of genes encoding potential anti-CRISPR/Cas9 or other DNA-modifying or -repair proteins. Second, pCrispr plasmids should carry genes encoding known suppressors of the mating-induced SOS response, e.g., *psiB* and *ssb,* which exist naturally on F and other large conjugative plasmids (63, 66–68). Finally, and perhaps most importantly, pCrispr/Cas9 elements as well as genes for surface-dispayed Nb’s should be introduced directly onto the conjugative plasmid to achieve a proliferative killing response among a targeted population.

Bacterial minicells are increasingly being deployed as factories for generation of biomolecules (69) and as vectors for delivery of cancer therapeutics (4–7). These applications have capitalized on the findings that minicells are readily engineered to function as biocatalysts and can readily be packaged with therapeutic molecules, e.g., siRNAs, shRNA plasmids, toxins, without appreciable drug leakage. Here, we showed that minicells naturally package functional T4SSs, plasmid cargoes, and Nb’s during aberrant cell division, with the important outcome that pCrispr plasmids are efficiently mobilized and robustly kill intact *E. coli* transconjugants. Such nonviable, conjugative vectors are especially well-suited for deployment as antimicrobials in various infection settings. We envision that minicells generated from probiotic strains, such as *E. coli* strain Nissle 1917, will prove ideal for antimicrobial intervention for the same reasons that Nissle 1917 minicells are increasingly being tested for cancer therapy. Specifically, these cells lack virulence factors, efficiently target hypoxic regions, reduce inflammation through interactions with the adaptive immune system, and are readily cleared (70, 71).

Disappointingly, Nb-producing *E. coli* minicells abundantly bound and induced aggregation of Ag-displaying *E. coli* and *P. aeruginosa* target cells, yet Nb/Ag pairing did not elevate minicell-mediated plasmid transfer or killing beyond frequencies achieved with unpaired matings (**Figs 4**, **5**). To reconcile these findings, we suggest that features such as small sizes and rounded architectures of minicells, which can prove beneficial for deployment in infection or cancer tumor environments, may also be problematic for targeted conjugative transfer. If surface-displayed Nb’s and T4SS nanomachines are spatially separated on the cell surface, for example, the small sizes and rounded architectures of minicells may impede simultaneous formation of Nb/Ag pairs and productive contacts between the T4SS and target cell (**Fig. 4B, schematic**). On the other hand, if Nb’s and T4SSs are closely juxtaposed on minicell envelopes, the Nb/Ag interaction could physically disrupt T4SS machine function or establishment of productive mating pairs. Various modifications in the Nb/Ag pairing system might achieve the goal of minicell-mediated targeted transfer; these could range from simple modulation of Nb or Ag production levels to adjustments in spacer lengths between the autotransporter OM β-barrel and Nb/Ag passenger domains to deployment of completely different types of surface ligands capable of binding receptors naturally displayed by target cells.

In summary, our findings indicate that bacterial PDSs composed of ligand-displaying cells or minicells and different conjugation systems offer considerable promise as an adjunct or alternative to antibiotics for antimicrobial intervention.

## MATERIALS AND METHODS

### Bacterial strains, plasmids, oligonucleotides and growth conditions

*E. coli* and *P. aeruginosa* strains used in this study are listed in Table S1. Plasmids and oligonucleotides used in this study are listed in Tables S1 and S2. Details of strain growth conditions and plasmid constructions are presented in File S1.

### Conjugation assays

Donor and recipient strains were grown overnight with antibiotic selections, diluted 100-fold in LB broth without antibiotics and incubated for 1.5 h at 37°C with shaking. When necessary, anhydrotetracycline (100 ng/ml final conc.; Sigma), sodium salicylate (1 μM final conc.; VWR), or arabinose (0.2% final conc.; GoldBio) was added for induction of gene expression, and cells were incubated for an additional 1 h at 37°C with shaking. Donors and recipients were mixed in a 1:1 ratio, determined by CFU counts, unless otherwise stated. For solid surface conjugation, 30 μl of the mating mix was added to sterile nitrocellulose filters (0.45 μm pore size; MF Millipore) on LB plates with inducers for gene expression when necessary. Plates were incubated at 37°C for the times specified, filters were resuspended in 1 ml fresh LB. For liquid conjugation, 50 μl of donor and recipient cells grown as described above were inoculated into 1 ml LB, and mating mixtures were incubated at 37°C for times specified. For minicell-mediated conjugation, minicells were mixed in a 10:1 ratio (determined by microscopic counts) with intact recipient cells. Matings were carried out in LB liquid media or on sterile nitrocellulose filters (0.025 μm pore size; MF Millipore) on LB plates at 37°C for times specified. Mating mixtures were vortexed and maintained on ice to terminate conjugation, and then serially diluted and spread on LB plates with antibiotics selective for donors (D), recipients (R), or transconjugants (Tc’s). The frequency of transfer was reported as the number of transconjugants per donor (Tc’s/D) or Tc’s/R for comparisons of transfer to different recipients in mixed matings or minicell matings. For conjugation-mediated killing assays, results were reported as the fold-decrease in viable transconjugants (Tc’s) resulting from transfer of the pCrispr targeting plasmids relative to transconjugants arising from transfer of the gRNA-control plasmid (normalized to 1). Minicell matings were routinely spread on LB plates selective for donor cells to further ensure the absence of viable donor cells during the mating period.

### Minicell purification

Minicells were purified from the minicell producing strain WM3886 essentially as described previously (72). WM3886 strains harboring plasmids of interest were grown overnight with appropriate antibiotic selections (Suppl. Materials and Methods), diluted 100-fold in 50 ml of LB broth without antibiotics and incubated for 1 h at 37° C with shaking. When necessary, anhydrotetracycline (10 ng/ml final concentration) was added to induce expression of genes encoding the Nb or Ag autotransporter fusions. Induced cell cultures were grown for 4 h at 37°C with shaking, and intact cells were removed by three rounds of differential centrifugation at 2,500 *x g* for 5 min. Minicells were pelleted by centrifugation at 10,000 *x g* for 15 min and resuspended in 0.5 ml of LB broth. Minicells were incubated at 37°C for 1 h after which ceftriaxone (100 mg/ml; Sigma) was added to kill growing intact cells. Cell debris was removed by centrifugation at 400 *x g*, and the supernatant was centrifuged at 10,000 *x g* to pellet minicells. Minicell purity was assessed by DIC microscopy and by plating of minicells on LB media to confirm the absence of cell growth. Resulting minicell preparations (∼10^10^ minicells/ml) were serially diluted for use in cell binding and conjugation assays.

### Protein detection by western blotting

Strains producing Strep-tagged TraN or TraN deleted of the N-proximal extracellular domain (TraNΔED) were grown overnight with antibiotic selections, diluted 100-fold in LB broth and incubated for 1.5 h at 37°C with shaking. Arabinose (0.2 % final concentration) was added for induction of *traN* or *traNΔED* gene expression, and cells were incubated for 1 h at 37°C with shaking. Cells (1 ml cultures) were pelleted by centrifugation at 5,000 *x g* for 5 min and resuspended in 100 μl of LB broth. An equivalent volume of 2×Laemmli buffer was added and the samples were boiled for 5 min prior to electrophoresis through SDS-12.5% polyacrylamide (30:0.8 acrylamide/bis-acrylamide) gels. Electrophoresed material was transferred to nitrocellulose membranes and blots were developed with primary antibodies against the Strep-tag (Genscript) or RNA polymerase (RNAP; BioLegend) and horseradish peroxidase (HRP)-conjugated secondary antibodies (BioRad) for detection by chemiluminescence.

### Visualization of Nb/Ag-mediated binding

Effects of Nb/Ag-mediated aggregation of intact cells or on minicell binding to intact cells were assessed visually and by microscopy. To assess effects on cellular aggregation, Nb- and Ag- (or null) strains were grown overnight with antibiotic selections, diluted 100-fold in LB broth without antibiotics and incubated for 1.5h at 37 °C with shaking. Strains were mixed in a 1:1 ratio, anhydrotetracycline (100 ng/ml final concentration) was added to induce Nb, Ag, or null production, mixtures were incubated for 1h at 37 °C with shaking, and then incubated without shaking for 2h at room temperature. Minicells purified from strain WM3886 engineered for Nb surface display were mixed with intact Ag- or null-producing *E. coli* MC4100 or *P. aeruginosa* PAO-1 cells, or with Ag- or null-producing *E. coli* cells additionally producing mCherry or mCerulean3, respectively. Minicell - cell mixtures (10 μl) were spotted on a 1% agarose pad and imaged with a Nikon A1 confocal microscope equipped with a Plan Apo 100x objective lens (Oil Immersion) equipped with mCherry hr, cfp hr, and DIC filters and NIS-Elements AR software.

### Statistical analyses

Conjugation experiments were performed at least three times in triplicate, and results were reported as average transfer frequencies with standard deviations shown as error bars. *p*-values between indicated data sets were calculated by the homoscedastic Student’s *t*-test. * p<0.05, ** p<0.001, *** p<0.0001.

### Data availability

The authors declare that all data supporting the findings of this study are available within the paper and its supplementary information files.

## Author Contributions

P.J.C. and Y.G.L. designed the research; Y.G.L, K.K., and N.O-K. performed the research; P.J.C, Y.G.L, K.K., and N.O-K. analyzed the data; P.J.C. and Y.G.L. wrote the paper with contributions from K.K. and N.O-K.

## ACKNOWLEDGMENTS

This work was supported by NIH Grant R35GM131892 to PJC. We thank members of the Christie lab for helpful discussions. We thank Dr. Joseph Mougous for strains and helpful discussions. We are grateful to Dr. Bill Margolin for strains and advice on minicell purification, Dr. Anna Konovalova for strains and helpful discussions, and Dr. Pratick Khara for helpful discussions.

## Legend for Supplementary Materials

**Table S1.** Strains and plasmids used in this study.

**Table S2.** Oligonucleotides used in this study.

**Suppl. File 1.** Supplemental Materials and Methods.

**Fig. S1.** Optimization of Nb/Ag pairing for pKM101 transfer.

**Fig. S2.** Optimization of Nb/Ag pairing for F and RP4-mediated transfer.

**Fig. S3.** Optimization of Nb/Ag pairing for DNA transfer from *E. coli* to *P. aeruginosa*.

